# Atherosclerotic plaque iron accumulation characterizes a distinct phase of intra-plaque haemorrhage and is associated with inflammation and remodelling

**DOI:** 10.64898/2026.04.27.721220

**Authors:** Elias B. Wieland, Aniek Plug, Benjamin Balluff, Marion Gijbels, Lele Han, Bryn Flinders, Eva Cuypers, Laura Kempen, Xioafei Li, Barend Mees, Erik Biessen, Marjo Donners, Pieter Goossens

**Affiliations:** Department of Pathology, Cardiovascular Research Institute Maastricht (CARIM), Maastricht University, Maastricht, The Netherlands; Department of Pathology, Maastricht University Medical Center (MUMC+), Maastricht, The Netherlands; M4I, The Maastricht MultiModal Molecular Imaging Institute, Maastricht University, Maastricht, The Netherlands; Laboratory of Immunology and Vaccinology, Faculty of Veterinary Medicine, FARAH, ULiège, Liège, Belgium; Laboratory of Immunophysiology, GIGA Institute, Liège University, Liège, Belgium; Department of Surgery, Maastricht University Medical Center (UMC)+, Maastricht, The Netherlands; Institute for Molecular Cardiovascular Research (IMCAR), RWTH Aachen University, Aachen, Germany

## Abstract

Intraplaque haemorrhage (IPH) is a hallmark of advanced atherosclerosis and a major risk factor for ischemic stroke and myocardial infarction. Current IPH classification focusses on extravascular erythrocyte presence as a proxy of acute bleeding, where iron detection generally indicates older haemorrhages. While often used interchangeably, a comprehensive analysis of transcriptional and metabolic context and impact of iron and erythrocyte deposition on the plaque is still lacking. Here, we investigate iron as a late-stage IPH hallmark in human atherosclerotic plaques.

We analysed erythrocyte-rich, iron-rich, and non-IPH regions in human carotid endarterectomy plaques by re-analysing a published transcriptomic dataset of 43 patient samples. In addition, we performed histological and immune phenotyping to define plaque traits associated with iron versus erythrocyte accumulation. Finally, we performed spatial metabolic profiling to functionally define iron-rich regions.

Although iron and erythrocyte deposits frequently co-localised, both co-related with different histological traits. While iron- and erythrocyte-rich regions shared transcriptomic features of advanced plaques compared with non-IPH regions, direct comparison showed differences in gene expression profiles. Iron deposition was associated with increased myeloid cell accumulation and a unique spatial metabolic signature distinct from erythrocyte-rich and non-IPH regions.

While sharing many characteristics with IPH plaques, the molecular, cellular and metabolic landscape of iron-rich regions is marked by features of plaque remodelling and repair. This makes iron deposition a unique hallmark of late-stage IPH, extending the current erythrocyte-based definition of IPH.

## Introduction

Atherosclerosis is a chronic inflammatory disease of large arteries, characterised by excessive lipid deposition forming lesions in the vessels’ intima layer (1). They can further develop from asymptomatic plaques into advanced, so-called unstable lesions. An important hallmark of advanced lesion instability is the presence of blood in the plaque, also termed intraplaque haemorrhage (IPH). Current understanding identifies leaky neovascularisation as the major source of IPH (2), and it is closely linked to several pathophysiological processes which are involved in the progression of atherosclerosis, such as hypoxia, angiogenesis, necrotic core formation, the local accumulation of macrophages, and the transition of endothelial cells to mesenchymal cells (EndMT) (3-6).

After the infiltration of blood in the atherosclerotic lesion, erythrocytes begin to degrade and release haemoglobin, which forms a complex with haptoglobin. Among the different macrophage subsets present in the plaque, a distinct population expressing CD163, the haemoglobin-haptoglobin scavenger receptor, is involved in the clearance of these bleeding sites (3, 7-9). Within the lysosomes of these macrophages, haemoglobin is degraded into heme and globin, after which heme is reduced by heme-oxygenase (HO-1) into free iron, which is then transported out of the macrophage via the iron receptor ferroportin. Hence, iron deposits in the plaque represent an end-product of metabolisation of IPH, further acting as a stimulator of macrophage function and phenotype (10, 11). Interestingly, both inflammatory and anti-inflammatory macrophages were shown to take up and metabolise iron (12-14). This calls for a phenotypic re-evaluation of myeloid cells that accumulate within areas of iron deposition in the plaque, helping to better understand how bleeding and especially its resolution influences macrophage populations and thus the status of local plaque inflammation.

The presence of IPH in plaque is mostly detected via magnetic resonance imaging (MRI) (15, 16). In *ex-vivo* plaque tissue, the current method of choice for IPH detection relies on a trained pathologist’s identification of erythrocyte accumulation (17). Even though these methods have proven efficient to predict the occurrence of adverse events (18), they do not account for older IPH events in plaques that, even when cleared by macrophages, may have led to a durable change in the plaque environment. To visualise these resolved bleeding events in the plaque, Perls’ iron staining offers a reliable method to visualise iron deposits (19). It is routinely used in other disease contexts such as the myelodysplastic syndrome or hemochromatosis (20-22).

While the association of IPH with pathophysiological processes is well established, the distinct impact of later-stage IPH resolution, particularly iron accumulation, on macrophage phenotypes and plaque characteristics remains unclear. Current definitions of IPH primarily rely on erythrocyte detection, but the unique molecular, cellular, and metabolic consequences of iron deposition as a late-stage hallmark have not been systematically explored. To address this gap, we characterized plaque regions with iron deposits using a multi-omics approach, comparing their metabolic, transcriptional, and macrophage profiles to erythrocyte-rich and non-IPH regions. By integrating transcriptomic proteomic and metabolomic analyses of human carotid endarterectomy plaques, we aimed to define the unique signatures associated with iron accumulation.

## Material and Methods

### Ethics statement and Patient Consent

All atherosclerotic plaque carotid endarterectomy patients’ samples were obtained in the context of the Maastricht Pathology Tissue Collection in line with the Dutch Code for Proper Secondary Use of Human Tissue (https://www.federa.org) and the local medical ethical committee (protocol number 16-4-181). This study conforms to the Declaration of Helsinki, all participants have given informed written consent prior to the inclusion.

### Human tissue procurement and processing

Human atherosclerotic plaques were obtained during carotid endarterectomy interventions. They were cut into parallel; transverse 5-mm-thick segments and alternating segments were snap-frozen in liquid nitrogen and stored at −80 °C while their flanking segments were fixed for 24 hours in formalin, decalcified for 4 hours before processing, and embedded in paraffin for histological evaluation. Plaque classification was performed on haematoxylin and eosin (H&E)–stained 4-µm-thick sections. Segments were stratified into thick fibrous cap atheromas and IPH containing plaques in accordance with the guidelines of Virmani *et al*. (17).

### Perls’ i ron staining

Perls’ iron staining (19) was performed on formalin fixed, paraffin embedded (FFPE)tissue sections and fresh frozen sections. A working solution was freshly prepared by mixing equal parts of 2% (w/v) potassium hexacyanoferrate (II) and 5.4% (v/v) concentrated hydrochloric acid in distilled water. Following deparaffinisation or thawing and fixation, sections were incubated with the filtered working solution for 30 minutes at room temperature and counterstained with Nuclear Fast Red (Sigma-Aldrich) for 5 minutes. Sections were mounted and imaged using the Pannoramic 1000 Digital scanner (3DHistech). Iron quantification was performed using QuPath software (v0.5.1) by automatic stain detection and normalisation to total tissue area (23). To determine the percentage of iron-positive areas per segment, the mean value of the two flanking FFPE sections was calculated and used for downstream analysis. Baseline characteristics of the cohort can be found in Table S1.

#### Mass spectrometry imaging

For mass spectrometry analysis, snap-frozen plaque tissue (n=3) was mounted using water and cut into 5 µm thick sections onto indium-tin oxide (ITO) coated slides as described earlier (24, 25).

The ITO coated slides were dried in a vacuum desiccator for 10 minutes, and several fiducial markers (Tipp-Ex, BIC, France) were applied before matrix deposition. Eight layers of 10 mg/mL α-Cyano-4-hydroxycinnamic acid (CHCA) (Sigma-Aldrich, Germany) in 70:30 (v/v) methanol:water were applied using the SunCollect pneumatic sprayer (SunChrom GmbH, Friedrichsdorf, Germany), as described previously (26). Matrix-assisted laser desorption/ionisation (MALDI)– Mass Spectrometry Imaging (MSI) was performed with a 9.4T Fourier transform ion cyclotron resonance mass spectrometer (Solarix; Bruker Daltonik GmbH, Bremen, Germany). The MALDI source is equipped with a Smartbeam II Nd:YAG UV laser. The measurements were performed with the laser operating at 2 kHz and by averaging 250 shots per pixel. The laser power was optimised to maximise the signal-to-noise ratio, which was at 70%. An internal calibration of the mass spectra was performed during acquisition using FTMS Processing V2.1 (Bruker Daltonik) with a maximum peak assignment tolerance of 10 ppm on the m/z scale. Data were acquired in magnitude mode and positive ion polarity at 50 µm lateral resolution in the mass range of m/z 40 to 800 using 1 million data points per spectrum. After acquisition, the ITO coated slides were washed with 70% methanol to remove matrix before being stained with Perls’ iron staining and counterstained with Nuclear Fast Red.

Slides were subsequently scanned (20×) using a Pannoramic P1000 Digital Scanner (3D HISTECH) to obtain digital high-resolution histological images, which have been co-registered to the MSI data using flexImaging 4.1 (Bruker Daltonik). A vascular pathologist, blinded to the MSI data, identified and digitally (Qupath v0.5.1) annotated size-matched areas of erythrocyte deposits, iron deposits, or regions without both features. The MSI data and the co-registered histology images with the annotated regions were imported to SCiLS Lab 2025a (Bruker Daltonik) for data analysis. From 1945 initial m/z features, 618 could be annotated using the Metaboscape (Bruker Daltonics) plugin in SCiLS Lab. For name matching of the m/z features, the human metabolome database (hmdb; v4.0) with a mass window of 3 ppm was used.

### Mass spectrometry imaging data analysis

The average non-normalised intensities for every metabolite, region and patient were exported to Rstudio (Version 2025.05.01+513). One-vs-One receiver-operating-characteristic (ROC) analyses were performed using the pROC (v1.19.0) package (27). For each metabolite, average non-normalised values per region and patient sample were used for the three pairwise comparisons (Non-IPH vs Iron, Non-IPH vs erythrocyte, Iron vs erythrocyte). The area under the curve (AUC) represented the effect size and one isoform per metabolite comparison was retained by keeping the entry with the furthest AUC from 0.5. Metabolites were classified as enriched in their respective histological region when the AUC was < 0.30 or > 0.70. Visualisations were produced with ggplot2 (v*3*.*5*.*2)* (28). KEGG over-representation analyses was conducted using KEGGREST (v1.5.0) *(29)*. Additional 3-group comparisons were done using the Kruskal–Wallis H test with p<0.05 regarded as statistically significant.

### Gene microarray data preparation

Gene microarray data of 43 plaque segments from 22 patients (the Maas-HPS study, GSE163154, previously described in detail (30, 31)), were used for this study.

Data were analysed using the R Bioconductor lumi package (v2.38.0) (32, 33). First, a variance stabilising transformation incorporated in the lumi package was performed (32). Then, the robust spline normalisation algorithm in the package was applied to normalise the data. Multiple transcript isoforms were mapped to the same gene based on the HUGO Gene Nomenclature Committee symbols by selecting the highest expressed transcript. For Venn diagrams and the heatmaps, iron and erythrocyte positivity was determined at the median, meaning that samples with an iron or erythrocyte enrichment above the median were termed as either iron-rich or erythrocyte-rich. For other analysis, iron and erythrocyte accumulation were used as continuous features. Other plaque traits were assessed by immunohistochemistry, as described by Jin *et al*. (2021) (30).

### Differential expression analysis

Differential gene expression analysis between iron-rich, erythrocyte-rich and non-IPH samples was performed using the limma package (v3.42.2) (34). Benjamini-Hochberg correction of the p-values for multiple testing and control of the false discovery rate (FDR) was applied.

### Automated cyclic immunofluorescent imaging

FFPE atherosclerotic plaque tissue from the MaasHPS cohort was processed as described above. Sections of 4-µm were cut and 1 representative sample was selected for high-plex MACSima™ multiplex fluorescent imaging using the REAscreen™ Immuno-oncology kit (human, FFPE, v01, Miltenyi Biotech) (Table S2). The staining and imaging procedure was performed according to the manufacturer’s protocol as described previously (35). Imaging data of a selection of 24 immune-related markers, based on staining specificity and biological relevance, were analysed (Table S2). From each marker’s image, background signal detected just prior to staining was subtracted to correct for autofluorescence. In the MACS^®^ iQ View software (Miltenyi Biotec), cells were segmented based on DAPI nuclear staining, after which median fluorescence intensities across all markers were extracted to Python (v3.13.7), transformed, and log normalised. The resulting single-cell expression matrix was used for unsupervised k-medoid clustering (kmedoids) and visualised using Uniform Manifold Approximation and Projection (UMAP) with the *umap* package for dimensionality reduction (36, 37). The clusters were manually annotated (Fig. S2) and these annotations were projected on the UMAP visualisation.

To assess regional differences, iron staining was performed as described above on the same tissue section. The image was manually annotated by a vascular pathologist into an iron-positive and iron-negative region, allowing to compare immune cell cluster distributions between these microenvironments.

### Iterative immunofluorescent imaging

Immunofluorescent imaging and staining procedure was performed as previously described by Guillot *et al*. (38). In brief, FFPE plaque sections from the MaasHPS cohort (n=40) were subjected to deparaffinisation and low-pH heat-induced antigen retrieval (HIER). To minimise autofluorescence, slides were exposed to intense LED light for one hour in a custom-made bleaching box, while being submerged in bleaching buffer (PBS with 30% H_2_O_2_ and 1M NaOH) (24). Plaques then underwent a cyclic staining, imaging, and stripping procedure for three cycles, where each cycle involved staining for two antigens plus a DAPI nuclear staining. Primary antibodies against the following targets were used: CD163, Lyve-1, CD206, Galectin3, HLA-DR and CD11c (Table S3). Rabbit antibodies were amplified using the VectaFluorTM Excel Amplification kit coupled to Alexa Fluor 488 (Vector/Brunschwig, cat# DK-1488), while mouse and rat primary antibodies were directly exposed to secondary antibodies conjugated with Alexa Fluor 674 or 594 (Invitrogen; cat# 10226162 and A-21471, respectively). After each staining cycle, whole-tissue imaging was conducted using a Pannoramic 250 digital slide scanner (3DHistech) at 20x magnification, with consistent exposure times within each cycle per channel (i.e., DAPI, FITC and TRITC). After the final cycle, Perl’s Prussian blue iron staining was performed on the same plaques, as described above, and slides were scanned on a Pannoramic 1000 digital slide scanner (3DHistech) at 20x magnification.

Image analysis was performed in QuPath (v0.5.1) (23) as follows: plaques were segmented into iron-containing regions and regions without iron based on the Perls’ iron staining. After manual removal of artefacts, the Stardist extension (v0.4.0) was used for cell segmentation based on DAPI nuclear staining, with consistent parameters for each plaque, including a 2 µm nuclear expansion (39). After cell quantification, cell positivity data were exported into Microsoft Excel (Microsoft, 2021) for data normalisation to total number of cells and annotation area. Prism 9 (GraphPad) was used for plotting and statistical analysis by non-parametric Wilcoxon paired tests.

## Results

### Erythrocyte and iron deposits co-localise in a majority of plaque samples but correlate with different plaque phenotypic traits

Current IPH detection by erythrocyte presence overlooks resolved or metabolised IPH events that can leave metabolic and inflammatory footprints within the plaque (2, 17). While iron accumulation represents the end-product of haemoglobin clearance, the plaque characteristics, myeloid phenotypes, and metabolic profiles associated with iron-rich regions remain undefined.

To identify the distribution of iron deposits relative to erythrocyte accumulation in the plaque, we first quantified iron and erythrocyte accumulation in the atherosclerotic plaque samples from the MaasHPS cohort (n=43) (Fig. 1A). Most plaques that contained erythrocyte accumulation also had iron deposits (n=15), and both often colocalise at the same sites within the lesion. However, erythrocyte deposits and iron deposits can be found alone as well, likely representing sites of early (initiation) and late (resolution) bleeding. Only few samples were exclusively iron-rich or erythrocyte-rich (n=4; n=5, respectively). We found that erythrocyte accumulation correlated with plaque calcification (*r*=0.34, p=0.03), plaque size (*r*=0.76, p<0.0001), lymph vessel density (*r*=0.66, p=0.0001), iNOS^+^ macrophages (*r*=0.51, p=0.004), PDGFRα^+^ fibroblasts (*r*=0.59, p<0.0001). Further, erythrocyte accumulation negatively correlated with collagen content (*r*=-0.33, p=0.04). Iron accumulation correlated positively with plaque size (*r*=0.54, p=0.0002), lymph vessel density (*r*=0.66, p<0.0001), iNOS^+^ macrophages (*r*=0.39, p=0.02), PDGFRα^+^ fibroblasts (*r*=0.69, p<0.0001) and CD3^+^ T-cells (*r*=0.32, p=0.04). In contrast to erythrocyte correlations, iron accumulation strongly negatively correlated with ARG1^+^ macrophages (*r*=-0.48, p=0.006) (Fig. 1B-C).

**Fig. 1.**
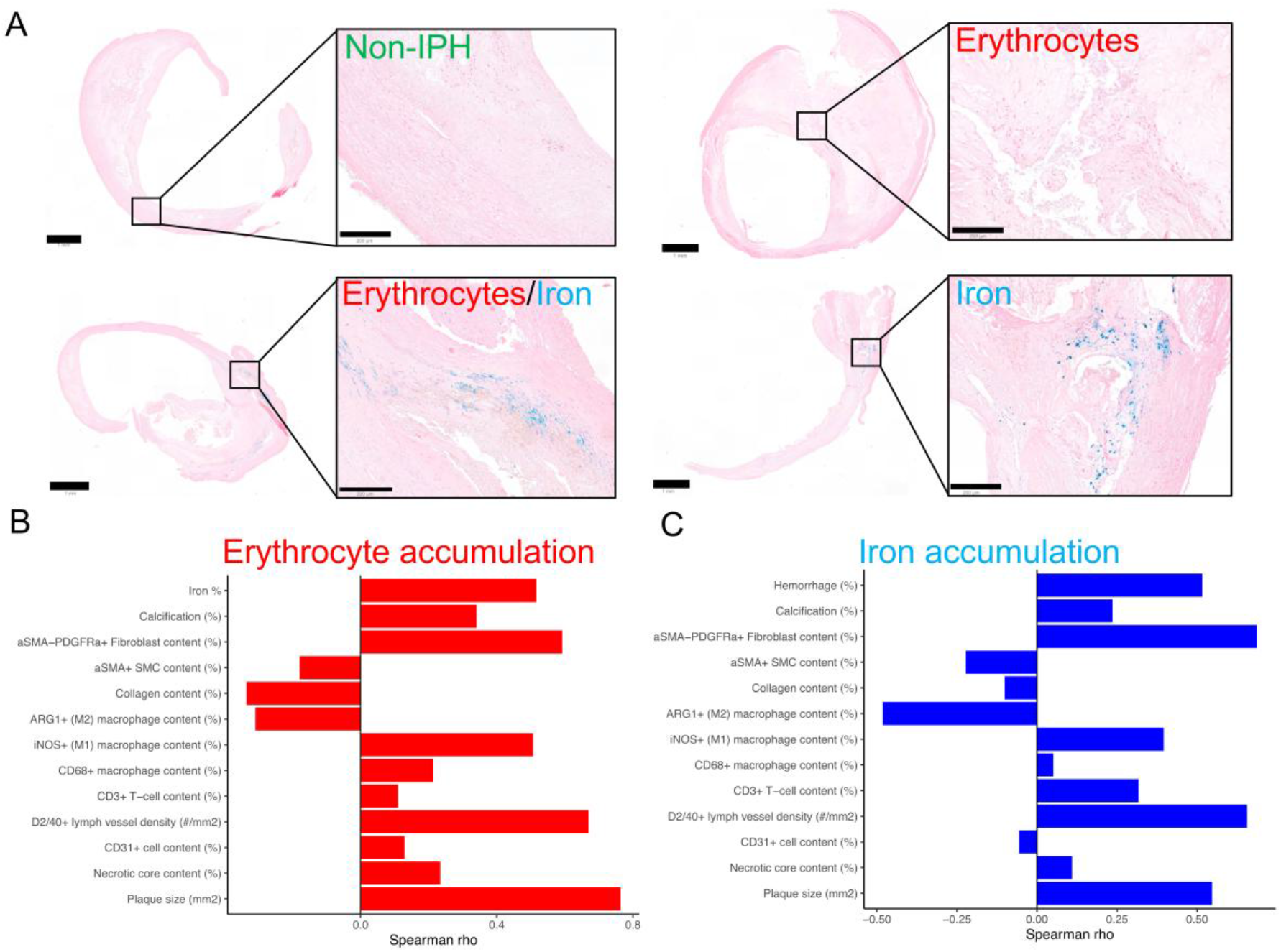
Distribution of iron within atherosclerotic plaques and its associations with plaque traits. **(A)** Representative images of iron deposits,presence of erythrocytes, the combination or the absence of both in human plaque samples (n=43).(**B-C)** Pearson correlation analysis of Irondeposition(%)(C) or Erythrocyte accumulation (%) (D) against plaque phenotypic traits.

### Iron and erythrocyte deposits share substantial transcriptomic overlap relative to non-IPH, yet retain distinct gene signatures

To further characterise differences in IPH and iron deposit, we re-analysed a bulk transcriptomic dataset, performed on segments flanking the stained sections mentioned above (30). First, we compared the transcriptomic profiles of all four groups, i.e. samples either being iron-rich, erythrocyte-rich, enriched in both or none (non-IPH). This revealed a clear separation between the transcriptomic profiles of samples containing both iron and erythrocytes and non-IPH samples (Fig 2A). The samples that were erythrocyte-rich clustered mainly with the samples that were enriched in both erythrocytes and iron deposit. Non-IPH samples were associated with expression of genes cells involved in tissue repair and mesenchymal cells (*NPNT, ACTC1, RGS5, CNN1, SRFP1, SOST, CYTL1*) (Fig 2A). In line with previous reports(40-44), plaques that contained both, erythrocytes and iron, were enriched in genes related to macrophage inflammatory functions, heme metabolism and foamy cells (*APOE, SPP1, CCL18, MMP7, MMP9, MMP12, MARCO, FABP4, VMO1, HMOX1* and *ADFP*) (Fig. 2A).

**Fig. 2.**
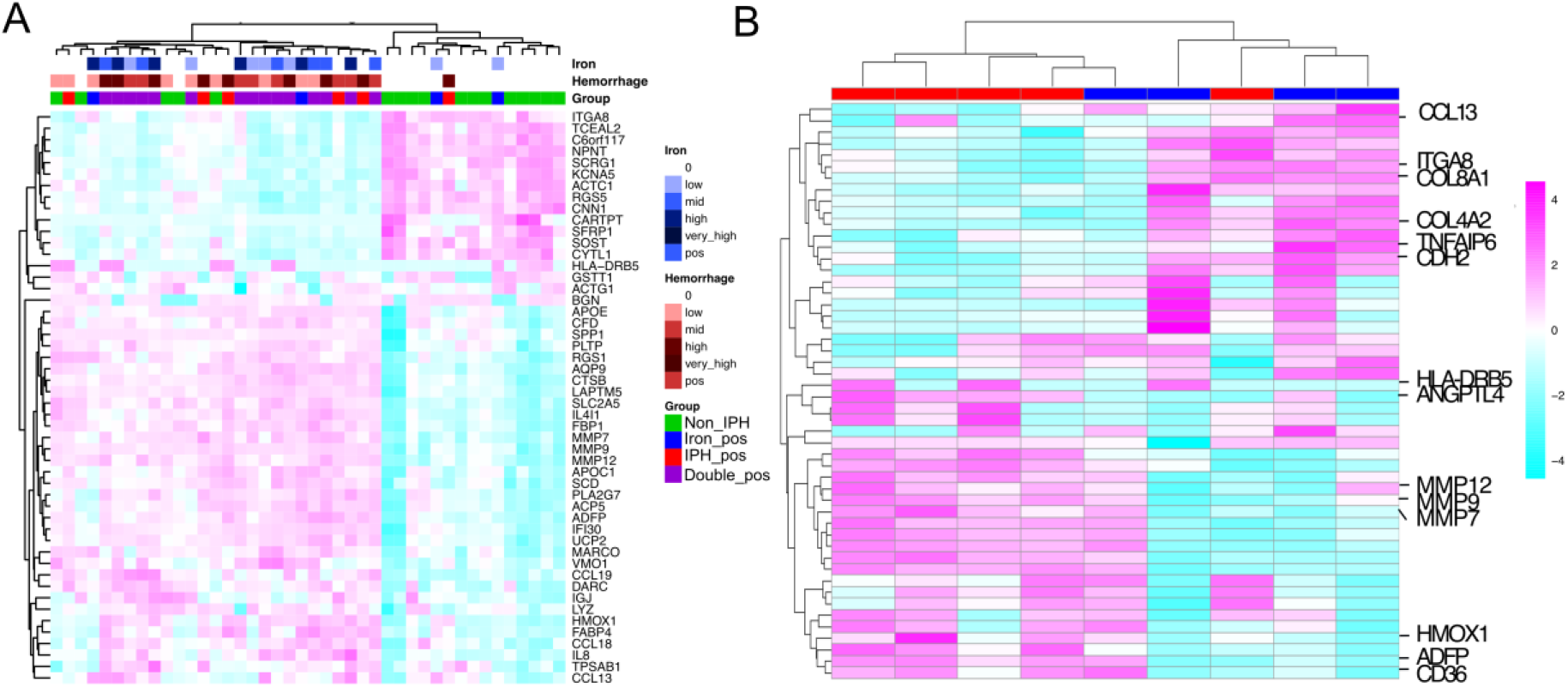
Association of transcriptomic profile and iron and erythrocyte content in atherosclerotic plaques. **(A)** Top 50 variable genes between plaque segments that are non-IPH, positive for iron, erythrocyte or positive for both. **(B)** Heatmap depicting the 50 most variable genes between erythrocyte-rich (n=5) and iron-rich plaque segments (n=4).

Next, we zoomed in on plaque segments that were only erythrocyte-rich or Iron-rich (Fig. 2B). Erythrocyte-rich samples showed mostly genes involved in lipid uptake and metabolism (*CD36, ADFP, ANGPTL4*), inflammation (*HLA-DRB5, MMP1, MMP9, MMP7*) and heme degradation (HMOX1) Among the genes that were associated with iron-rich samples, we observed CCL13, ITGA8, *COL8A1, COL4A2, TNFAIP6* and *CDH2*. These genes were reported to play a role in immune cell infiltration, but also in processes like endothelial-to-mesenchymal transition and tissue repair(45-49) (Fig. 2B).

### Multiplex Immunofluorescent imaging reveals co-localisation of immune cells with i ron-deposit in the plaque

As iron accumulation was associated with myeloid and lymphocyte cell populations and functions (Fig. 1B, Fig. 2), we performed exploratory automated cyclic multiplex immunofluorescence imaging to investigate the local cell landscape linked to iron deposits (Fig. 3A). In total, we detected 6198 cells and performed UMAP clustering using 24 athero-relevant cluster-defining markers (Fig. 3B; Table S2). By manual cell annotation based on recent work by others and us (50, 51), we defined 17 immune cell cluster (Fig. 3B; Fig. S1). Overall, we detected nine macrophage clusters, two CD8^+^ T cell, one mast cell clusters, and five other subsets (Fig. 3B). To decipher which subsets are spatially linked with iron accumulation in the plaque, we compared their abundance in areas segmented on iron staining positivity (Fig. 3C). Comparing the cellular composition of the iron and non-iron regions (3177 cells in the iron region, 3021 cells in the non-iron region), albeit based on a single plaque, a clear iron-specific cell profile could be identified (Fig. 3D). In line with their reported role in iron metabolism and uptake, CD163^+^ monocytes and macrophages were enriched within the iron deposition (7, 52). Further, Galectin-3 expressing macrophages and Lyve-1^+^ macrophages were found to be enriched within the iron deposition, while CD206^+^ macrophages and HLA-DR^+^ cells were higher in the non-IPH area of this specific plaque (Fig. 3E, Table S4).

**Fig. 3.**
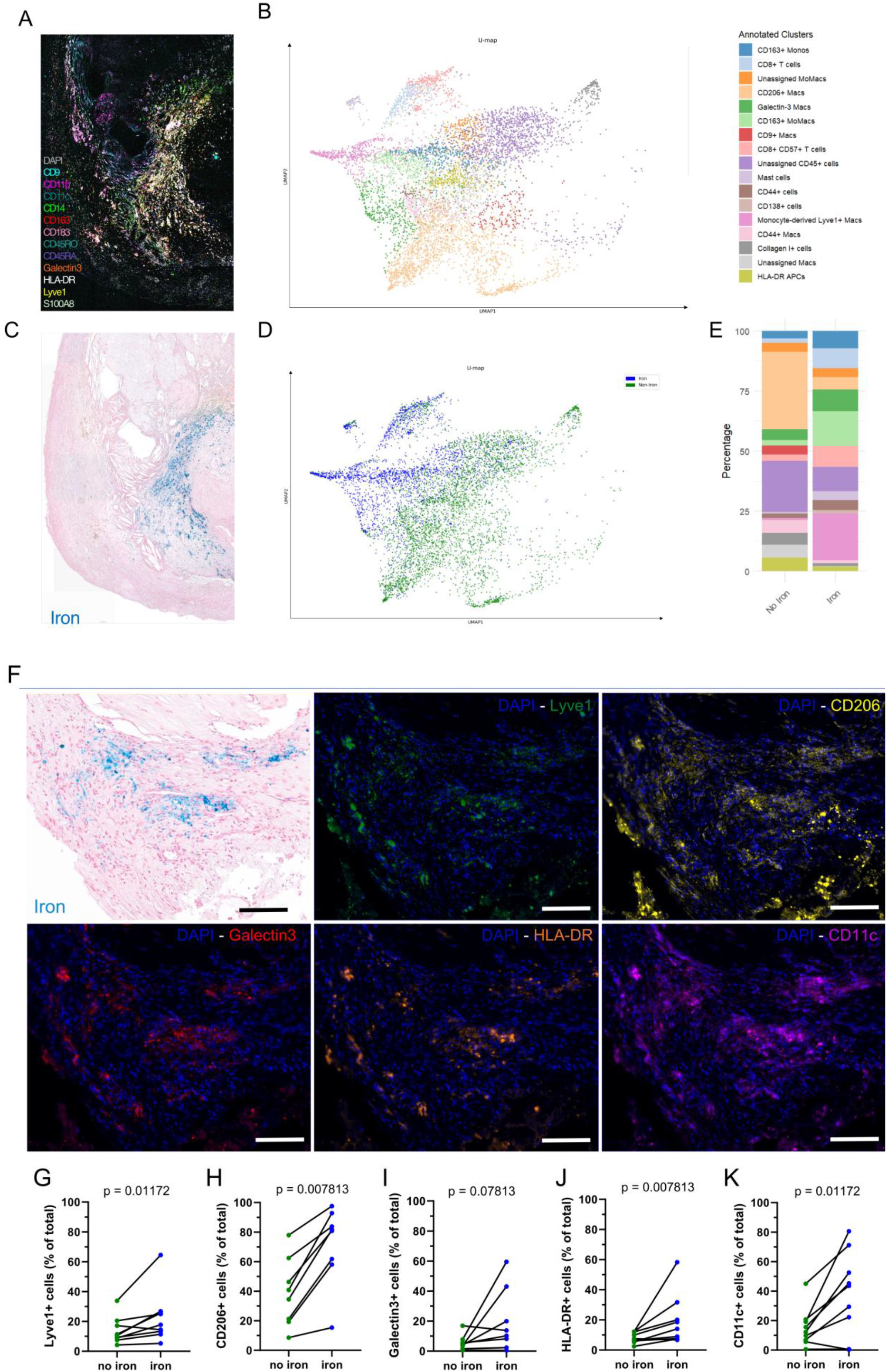
Spatial association of immune cells with iron accumulation. **(A)** Representative image of the 24-marker multiplex imaging experiment (n=1), visualizing 13 of the markers. **(B)** UMAP representation of all 17 detected cell subsets and their annotations. **(C)** Iron histochemistry image used to segment the area of iron accumulation. **(D)** UMAP representation with overlayed subsets enriched in iron (blue) and non-iron (green) areas. **(E)** Percentual comparison of cell populations between iron and non-iron areas. **(F)** Representative image of iron accumulation and 5 myeloid cell marker stainings. **(G-K)** Quantifications comparing Lyve-1^+^, CD206^+^, Galectin3^+^, HLA-DR^+^ and CD11c^+^ cells between iron and non-iron containing areas. Wilcoxon pairwise comparisons were used for statistics and scale bars indicate 100 µm.

To confirm the macrophage enrichment in areas with iron deposit in a larger set of plaques, we stained for Lyve-1, CD206, Galectin-3, HLA-DR and CD11c on plaque sections from the MaasHPS cohort (n=9) followed by a counterstaining for iron deposits (Fig. 3F). Interestingly, iron-rich areas displayed a higher cell density compared to regions without iron (Fig. S2). This suggests that regions with iron deposit are areas of high immune activity, consistent with the above-described conclusions (Fig. 1-2).

Despite some plaque heterogeneity, the staining showed that Lyve-1^+^ is indeed significantly enriched in the iron areas, but also^+^, CD206^+^, HLA-DR^+^ and CD11c^+^ cells were significantly enriched in these positive areas (Fig. 3G-K). Overall, these results show a relative enrichment of different myeloid subsets in the environment of iron deposit, compared to non-IPH regions (Fig. 3F-K).

### Regions of erythrocyte and iron accumulation are metabolically distinct from non-IPH regions and also differ from each other

To further characterise differences between the spatial niches that represent erythrocyte accumulation and iron deposition, mass spectrometry imaging (MSI) was performed on human carotid endarterectomy sections (n=3 patients). For each patient sample, size-matched annotations of these regions were made to compare their metabolomic profile in three independent receiver-operator curve (ROC) analyses (Fig. 4A-B; Supp. Information).

**Fig. 4.**
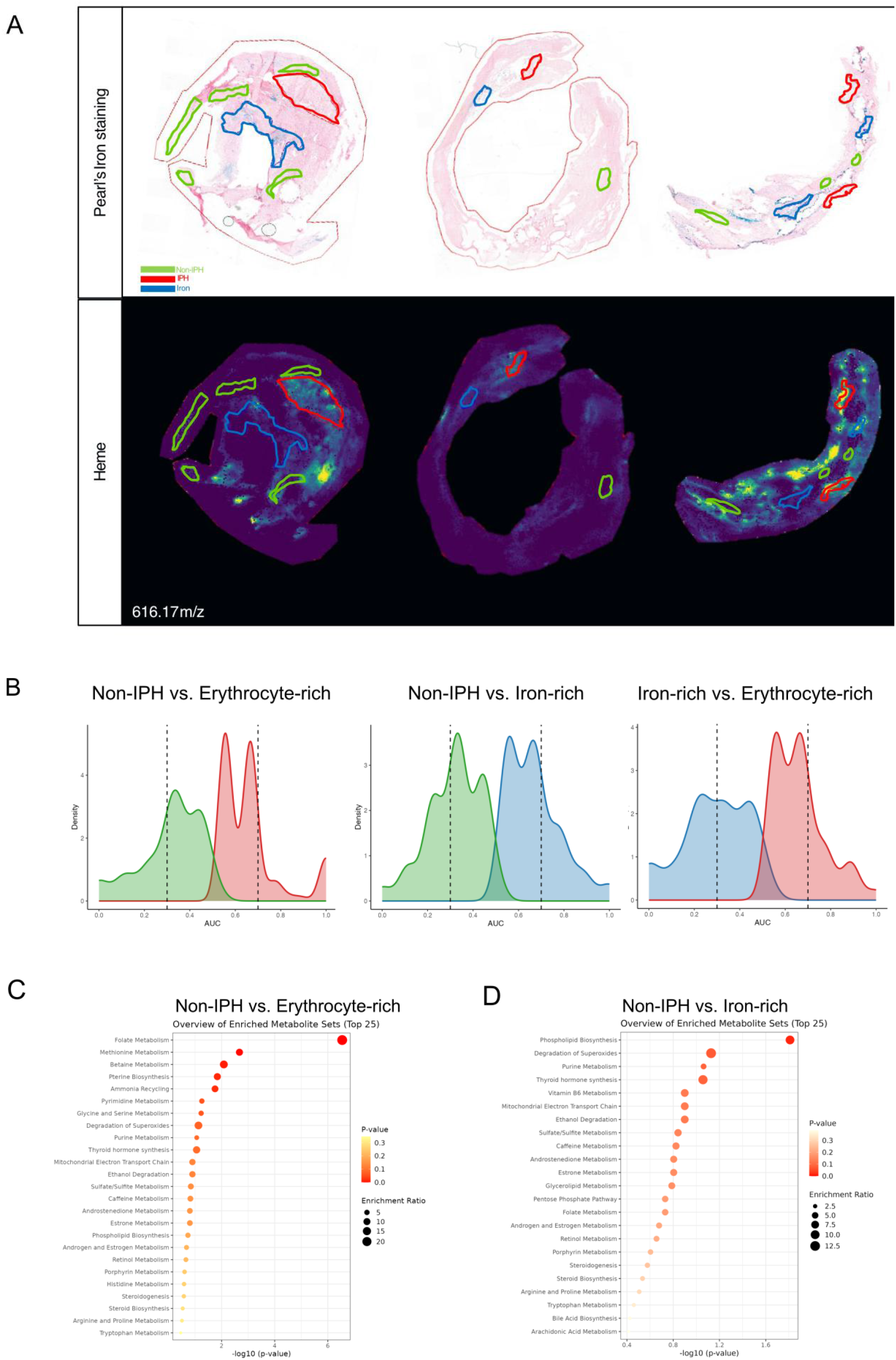
Spatial metabolomics to decipher regional metabolic differences between non-IPH, erythrocyte and iron regions. (A) Representative mass spectrometry image showing the iron histology and the histological annotations of non-IPH, erythrocyte and iron regions, and the distribution of hemein the plaque sections (n=3). (B) Density histograms visualising the ROC analyses comparing the three histological regions. Histograms show the distribution of metabolites that discriminate between the different regions. Metabolites below or above these thresholds were considered for functional analysis. The cut-off was set at 0.3 and 0.7 shown by the dashed line. (C-D) Pathway analysis of the different lists of discriminative metabolites per comparison.

When comparing non-IPH with erythrocyte-rich regions, we identified 123 metabolites specific to non-IPH regions and 34 metabolites specific to erythrocyte-rich regions. In the comparison between non-IPH and iron-rich regions, 110 and 67 metabolites, respectively, were found to be specific. Finally, direct comparison of iron-rich versus erythrocyte-rich regions revealed 138 metabolites enriched in iron-rich regions and 64 enriched in erythrocyte-rich regions (Fig. 4B; Supplementary Information). Confirming the histological annotations, heme, a major component of red blood cells, strongly discriminatederythrocyte-rich regions from both non-IPH and iron-rich regions, achieving an AUC = 1 in both comparisons (Fig. 4A; Supplementary Information).

To further interpret these metabolite-level differences, we conducted metabolic pathway enrichment analysis using MetaboAnalyst (v6.0) (53). Erythrocyte-rich regions were primarily enriched in pathways related to folate, methionine, betaine and pterin/pteridine metabolism, and ammonia recycling relative to non-IPH regions (Fig. 4C). Folate acid metabolism is reported to be important in activated macrophages (54). In contrast, iron-rich regions showed strong enrichment in phospholipid biosynthesis and superoxide degradation pathways (Fig. 4D).

Direct comparison of erythrocyte-rich and iron-rich regions also demonstrated distinct pathway enrichment patterns (Fig. S3A-B). Compared to iron-rich regions, erythrocyte-rich regions largely conserved the pathway enrichment observed in the non-IPH vs. erythrocyte-rich comparison and additionally exhibited enrichment in thyroid hormone synthesis (Fig. S3A). Finally, compared to erythrocyte-rich regions, iron-rich regions were enriched in purine metabolism, vitamin B6 metabolism, and phospholipid biosynthesis (Fig. S3B).

Together, these findings suggest that, iron-rich and erythrocyte-rich regions exhibit distinct local metabolic and cellular identities.

## Discussion

In this study, we demonstrate that iron accumulation in atherosclerotic plaques marks a biologically distinct microenvironment, characterised by a immune, cellular and metabolic profile that resembles intra-plaque haemorrhage, but markedly differs from the erythrocyte-rich regions that classically define IPH.

At the tissue level, we found that only limited amount of plaque segments contained uniquely erythrocyte- or iron-rich regions and that in most plaques erythrocyte accumulation and the deposition of iron coincide. We further observed that iron accumulation was linked to hallmarks of advanced and unstable plaques, correlating to blood and lymph vessel density, inflammatory macrophages, fibroblasts, and T cells. These findings suggest that iron-rich and erythrocyte-rich plaques strongly resemble each other in terms of general plaque morphology, probably because the time span of erythrocyte-accumulation until the appearance of iron deposits is relatively short, compared to the long-term process of atherogenesis. Further these findings suggest that differences between erythrocyte and iron accumulation are the easiest to detect by studying their direct micro-environment.

At the histological level, we identified several macrophage markers that were locally associated with iron-rich regions. Macrophages positive for CD206 were previously reported to co-localise with iron deposits in the atherosclerotic plaque (7), corroborating our findings. While Lyve-1^+^ and CD206^+^ macrophages are associated with an atheroprotective, pro-resolving phenotype (55-57), HLA-DR and CD11c have been associated with progressing inflammation in atherosclerosis (57, 58). The colocalization of these cells with iron deposits suggests that iron possesses a dual stimulatory role in atherosclerosis. Whereas heme and its degradation products such as iron were previously shown to directly stimulate inflammatory polarisation of macrophages (59), several studies showed that anti-inflammatory macrophages have potentially higher capacities in iron uptake than pro-inflammatory macrophages that might be more efficient in long term iron storage (11-14). Even if these studies were mostly conducted *in vitro*, potentially using different experimental parameters, our findings that iron deposits in human plaque co-localise with both anti- and pro-inflammatory macrophages seem to be in line with these observations.

At molecular level, we observed a high similarity between the transcriptomic profiles associated with erythrocyte and iron deposits compared to plaques without IPH. However, we noted differences between erythrocyte-rich and iron-rich plaques when comparing their reciprocally most variable genes, with several genes uniquely associated to iron-rich plaques that are involved in inflammatory cell recruitment (*CCL13, ITGA8*) (45, 60), inflammation resolution (*COL8A1, COL4A2, TNFAIP6*) and endothelial-to-mesenchymal transition (EndMT) (*ITGA8, COL8A1, CDH2*) (46, 61, 62). Interestingly, EndMT was recently described to be induced by IPH-associated CD163^+^ macrophages (6). We indeed observed presence relative abundance of CD163 myeloid cells within these regions of iron accumulation. This is in line with the hypothesis that the metabolism of erythrocytes by CD163^+^ macrophages triggers plaque remodelling and fibrosis (6, 63).

The local metabolic profiles of erythrocyte- and iron-rich regions are characterised by distinct pathways. We found that the most important erythrocyte-associated metabolic pathway was folate acid metabolism, which is reported to be a marker of activated macrophages (64). Moreover, folate receptor beta (FR-β) is used for *in vivo* imaging approaches to detect macrophage accumulation at inflammatory sites (65-67). This suggests the inflammatory nature of erythrocyte-rich regions. Iron-rich regions on the other hand are distinguished from non-IPH regions by an enrichment in phospholipid metabolism and the degradation of superoxides. As phospholipids are essential constituents of cell membranes, the enrichment in their metabolism could reflect local cell accumulation, in line with the increased cell density that we observed in areas of iron deposits (Fig. S3). The degradation of superoxides is reported to be a part of reparative processes associated with iron-rich regions in plaques (68). Iron-compared to erythrocyte-rich regions also showed an enrichment in purine and vitamin B6 metabolism. Purine metabolism is essential for DNA synthesis and hints, together with phospholipid accumulation, at a proliferative potential of cells within the iron-rich regions (69). Interestingly, vitamin B6 was reported to exert anti-inflammatory effects and inhibited atherosclerosis development in a rat atherosclerosis model (70-72). However, most of these studies focussed on functional effects of vitamin B6 supplementations, not necessarily reflecting physiological doses of vitamin B6 or investigating local effects of vitamin B6 metabolism.

Our findings indicate that iron accumulation marks a specific stage in the process of intraplaque haemorrhage, distinct from erythrocyte accumulation. The anti-inflammatory and tissue-remodelling features associated with iron accumulation, indicate that iron deposition represents the aftermath of erythrocyte accumulation. Thus, including iron detection in the characterisation of IPH might provide a more complete comprehension of atherosclerotic plaque samples and enables a more detailed stratification of patient samples.

The *in-situ* detection of iron deposition is not restricted to *ex vivo* histological stainings. In oncology, “ferronostics” describes the intravital detection of iron in a human xenograft glioma model using PET-CT and the experimental labile iron pool-sensing radiotracer [18F]TRX (73). Our findings suggest that similar iron-targeted imaging strategies could be adopted for atherosclerosis, though their application in cardiovascular disease has yet to be tested. Other modalities, such as T2*-weighted magnetic resonance imaging (MRI) and contrast-enhanced MRI, may also offer non-invasive approaches to detect intravital iron in plaques, although the latter is currently limited to visualising the localisation of injected small superparamagnetic iron oxide nanoparticles, instead of the endogenously present iron (16, 74).

More generally, the source and role of iron in cardiovascular disease have been subject to debate (75). The ‘iron hypothesis’, first described in the previous century, suggests that systemic iron overload contributes to the progression of ischemic heart disease (76). While injection of apolipoprotein E-deficient mice with iron led to its’ deposition in the plaque and was found to be associated with oxidative stress and endothelial dysfunction (77, 78), another study using *ApoE*-deficient mice found that experimental iron accumulation in macrophages had no effect on plaque lesion development (79). In humans, iron administration via the contrast agent ferumoxtran was associated with increased apoptosis within plaques (16), yet patients with hereditary hemochromatosis, who experience lifelong iron overload, do not exhibit a higher incidence of atherosclerosis, strongly challenging the importance of the iron hypothesis in atherogenesis (75).

Further studies in larger patient cohorts with a larger number of iron-rich plaques are required to further assess the clinical impact and relevance of iron deposit in human atherosclerosis. In conclusion, our findings support an expanded model of IPH that encompasses both the acute phase of erythrocyte accumulation and a subsequent iron accumulation phase, characterized by a distinct metabolic and cellular profile, that may be associated with linked to tissue remodelling and inflammation. By defining this secondary phase, we refine the traditional erythrocyte-centric definition of IPH. Consequently, incorporating iron detection may enhance plaque characterisation, improve patient and plaque sample stratification and guide the development of IPH imaging strategies.

## Acknowledgments

This work has been supported by the Dutch Heart Foundation (Dekker 2020T042, to P.G.) and EIC Horizon 2020 Pathfinder (ABCardionostics, to E.B. and P.G.). The authors would like to thank Miltenyi Biotec (Germany) and Evelien Van Hamme (VIB, Belgium) for the support with the multiplex imaging. Declaration of interest: none.

## References

1. Bjorkegren JLM, Lusis AJ. Atherosclerosis: Recent developments. Cell. 2022;185(10):1630–45.

2. Kodama T, Narula N, Agozzino M, Arbustini E. Pathology of plaque haemorrhage and neovascularization of coronary artery. J Cardiovasc Med (Hagerstown). 2012;13(10):620–7.

3. Balmos IA, Slevin M, Brinzaniuc K, Muresan AV, Suciu H, Molnár GB, et al. Intraplaque Neovascularization, CD68+ and iNOS2+ Macrophage Infiltrate Intensity Are Associated with Atherothrombosis and Intraplaque Hemorrhage in Severe Carotid Atherosclerosis. Biomedicines. 2023;11(12).

4. Guo L, Harari E, Virmani R, Finn AV. Linking Hemorrhage, Angiogenesis, Macrophages, and Iron Metabolism in Atherosclerotic Vascular Diseases. Arteriosclerosis, Thrombosis, and Vascular Biology. 2017;37(4):e33–e9.

5. Guo L, Akahori H, Harari E, Smith SL, Polavarapu R, Karmali V, et al. CD163+ macrophages promote angiogenesis and vascular permeability accompanied by inflammation in atherosclerosis. J Clin Invest. 2018;128(3):1106–24.

6. Mori M, Sakamoto A, Kawakami R, Guo L, Slenders L, Mosquera JV, et al. CD163+ Macrophages Induce Endothelial-to-Mesenchymal Transition in Atheroma. Circulation Research. 2024;135(2):e4–e23.

7. Finn AV, Nakano M, Polavarapu R, Karmali V, Saeed O, Zhao X, et al. Hemoglobin directs macrophage differentiation and prevents foam cell formation in human atherosclerotic plaques. J Am Coll Cardiol. 2012;59(2):166–77.

8. Nagy E, Eaton JW, Jeney V, Soares MP, Varga Z, Galajda Z, et al. Red cells, hemoglobin, heme, iron, and atherogenesis. Arterioscler Thromb Vasc Biol. 2010;30(7):1347–53.

9. Boyle JJ, Harrington HA, Piper E, Elderfield K, Stark J, Landis RC, et al. Coronary intraplaque hemorrhage evokes a novel atheroprotective macrophage phenotype. Am J Pathol. 2009;174(3):1097–108.

10. Cai J, Zhang M, Liu Y, Li H, Shang L, Xu T, et al. Iron accumulation in macrophages promotes the formation of foam cells and development of atherosclerosis. Cell Biosci. 2020;10(1):137.

11. Gan ZS, Wang QQ, Li JH, Wang XL, Wang YZ, D. HH. Iron Reduces M1 Macrophage Polarization in RAW264.7 Macrophages Associated with Inhibition of STAT1. Mediators Inflamm. 2017;2017:8570818.

12. Corna G, Campana L, Pignatti E, Castiglioni A, Tagliafico E, Bosurgi L, et al. Polarization dictates iron handling by inflammatory and alternatively activated macrophages. Haematologica. 2010;95(11):1814–22.

13. Recalcati S, Locati M, Marini A, Santambrogio P, Zaninotto F, De Pizzol M, et al. Differential regulation of iron homeostasis during human macrophage polarized activation. European Journal of Immunology. 2010;40(3):824–35.

14. Ludwig N, Cucinelli S, Hametner S, Muckenthaler MU, Schirmer L. Iron scavenging and myeloid cell polarization. Trends in Immunology. 2024;45(8):625–38.

15. Cappendijk VC, Cleutjens KB, Heeneman S, Schurink GW, Welten RJ, Kessels AG, et al. In vivo detection of hemorrhage in human atherosclerotic plaques with magnetic resonance imaging. J Magn Reson Imaging. 2004;20(1):105–10.

16. Segers FME, Ruder AV, Westra MM, Lammers T, Dadfar SM, Roemhild K, et al. Magnetic resonance imaging contrast-enhancement with superparamagnetic iron oxide nanoparticles amplifies macrophage foam cell apoptosis in human and murine atherosclerosis. Cardiovasc Res. 2023;118(17):3346–59.

17. Virmani R, Burke AP, Kolodgie FD, Farb A. Vulnerable Plaque: The Pathology of Unstable Coronary Lesions. Journal of Interventional Cardiology. 2002;15(6):439–46.

18. Bos D, Arshi B, van den Bouwhuijsen Qja, Ikram MK, Selwaness M, Vernooij MW, et al. Atherosclerotic Carotid Plaque Composition and Incident Stroke and Coronary Events. J Am Coll Cardiol. 2021;77(11):1426–35.

19. Sonoda A, Nihei M, Shinkawa N, Kakizaki E, Yukawa N. Perls’ Prussian blue staining and chemistry of Prussian blue and Turnbull blue. Forensic Science International: Synergy. 2025;11:100627.

20. Seo IS, Li C-Y, Yam LT. Myelodysplastic Syndrome: Diagnostic Implications of Cytochemical and Immunocytochemical Studies. Mayo Clinic Proceedings. 1993;68(1):47–53.

21. Rovai A, Chung B, Hu Q, Hook S, Yuan Q, Kempf T, et al. In vivo adenine base editing reverts C282Y and improves iron metabolism in hemochromatosis mice. Nature Communications. 2022;13(1):5215.

22. Lours C, Cottin L, Wiber M, Andrieu V, Baccini V, Baseggio L, et al. Perls’ Stain Guidelines from the French-Speaking Cellular Hematology Group (GFHC). Diagnostics (Basel). 2022;12(7).

23. Bankhead P, Loughrey MB, Fernández JA, Dombrowski Y, McArt DG, Dunne PD, et al. QuPath: Open source software for digital pathology image analysis. Scientific Reports. 2017;7(1):16878.

24. Wieland EB, Kempen LJAP, Lu C, Donners MMPC, Biessen EAL, Goossens P. Protocol for multispectral imaging on cryosections to map myeloid cell heterogeneity in its spatial context. STAR Protocols. 2023;4(4):102601.

25. Cao J, Martin-Lorenzo M, van Kuijk K, Wieland EB, Gijbels MJ, Claes BSR, et al. Spatial Metabolomics Identifies LPC(18:0) and LPA(18:1) in Advanced Atheroma With Translation to Plasma for Cardiovascular Risk Estimation. Arteriosclerosis, Thrombosis, and Vascular Biology. 2024;44(3):741–54.

26. Sun N, Ly A, Meding S, Witting M, Hauck SM, Ueffing M, et al. High-resolution metabolite imaging of light and dark treated retina using MALDI-FTICR mass spectrometry. Proteomics. 2014;14(7-8):913-23.

27. Robin X, Turck N, Hainard A, Tiberti N, Lisacek F, Sanchez J-C, et al. pROC: an open-source package for R and S+ to analyze and compare ROC curves. BMC Bioinformatics. 2011;12(1):77.

28. Wickham H. ggplot2: Elegant Graphics for Data Analysis. Springer-Verlag New York. 2016.

29. Tenenbaum D MB. KEGGREST: Client-side REST access to the Kyoto Encyclopedia of Genes and Genomes (KEGG). R package version 1500. 2025.

30. Jin H, Goossens P, Juhasz P, Eijgelaar W, Manca M, Karel JM, et al. Integrative multiomics analysis of human atherosclerosis reveals a serum response factor-driven network associated with intraplaque hemorrhage. Clinical and Translational Medicine. 2021;11(6):e458.

31. Jin H BE. Maastricht Human Plaque Study (MaasHPS). Gene Expression Omnibus. 2020.

32. Du P, Kibbe WA, Lin SM. lumi: a pipeline for processing Illumina microarray. Bioinformatics. 2008;24(13):1547–8.

33. The open-source data science company. 2022.

34. Ritchie ME, Phipson B, Wu D, Hu Y, Law CW, Shi W, et al. limma powers differential expression analyses for RNA-sequencing and microarray studies. Nucleic acids research. 2015;43(7):e47–e.

35. Kinkhabwala A, Herbel C, Pankratz J, Yushchenko DA, Rüberg S, Praveen P, et al. MACSima imaging cyclic staining (MICS) technology reveals combinatorial target pairs for CAR T cell treatment of solid tumors. Scientific Reports. 2022;12(1):1911.

36. Schubert E, Lenssen L. Fast k-medoids Clustering in Rust and Python. Journal of Open Source Software. 2022;7(75).

37. McInnes L, Healy J, Saul N, Großberger L. UMAP: Uniform Manifold Approximation and Projection. Journal of Open Source Software. 2018;3(29).

38. Guillot A, Kohlhepp MS, Bruneau A, Heymann F, Tacke F. Deciphering the Immune Microenvironment on A Single Archival Formalin-Fixed Paraffin-Embedded Tissue Section by An Immediately Implementable Multiplex Fluorescence Immunostaining Protocol. Cancers. 2020;12(9):2449.

39. Schmidt U, Weigert M, Broaddus C, Myers G. Cell Detection with Star-Convex Polygons. Medical Image Computing and Computer Assisted Intervention – MICCAI 2018: Springer International Publishing; 2018. p. 265–73.

40. Xue S, Tang H, Zhao G, Fang C, Shen Y, Yan D, et al. C–C motif ligand 8 promotes atherosclerosis via NADPH oxidase 2/reactive oxygen species-induced endothelial permeability increase. Free Radical Biology and Medicine. 2021;167:181–92.

41. Gough PJ, Gomez IG, Wille PT, Raines EW. Macrophage expression of active MMP-9 induces acute plaque disruption in apoE-deficient mice. J Clin Invest. 2006;116(1):59–69.

42. Grebe A, Hoss F, Latz E. NLRP3 Inflammasome and the IL-1 Pathway in Atherosclerosis. Circulation Research. 2018;122(12):1722–40.

43. Bengtsson E, Hultman K, Edsfeldt A, Persson A, Nitulescu M, Nilsson J, et al. CD163+ macrophages are associated with a vulnerable plaque phenotype in human carotid plaques. Sci Rep. 2020;10(1):14362.

44. Gissler MC, Mwinyella T, Morguet C, Heitlinger S, Li X, Horstmann H, et al. Macrophage expressed CD36 promotes plaque vulnerability in atherosclerosis. European Heart Journal. 2023;44(Supplement_2):ehad655.3080.

45. Mendez-Enriquez E, García-Zepeda EA. The multiple faces of CCL13 in immunity and inflammation. Inflammopharmacology. 2013;21(6):397–406.

46. Ryu J, Koh Y, Park H, Kim DY, Kim DC, Byun JM, et al. Highly Expressed Integrin-α8 Induces Epithelial to Mesenchymal Transition-Like Features in Multiple Myeloma with Early Relapse. Mol Cells. 2016;39(12):898–908.

47. Li Q, Ye L, Talapaneni S, Meng Y, Wang CR, Kalindjian K, et al. COL8A1 regulates endothelial phenotype in inflammatory endothelial-to-mesenchymal transition. Am J Physiol Heart Circ Physiol. 2025;329(5):H1331–h46.

48. Watanabe R, Watanabe H, Takahashi Y, Kojima M, Konii H, Watanabe K, et al. Atheroprotective Effects of Tumor Necrosis Factor–Stimulated Gene-6. JACC: Basic to Translational Science. 2016;1(6):494–509.

49. Welch-Reardon KM, Wu N, Hughes CCW. A Role for Partial Endothelial–Mesenchymal Transitions in Angiogenesis? Arteriosclerosis, Thrombosis, and Vascular Biology. 2015;35(2):303–8.

50. Vallejo J, Cochain C, Zernecke A, Ley K. Heterogeneity of immune cells in human atherosclerosis revealed by scRNA-Seq. Cardiovasc Res. 2021;117(13):2537–43.

51. Goossens P, Lu C, Cao J, Gijbels MJ, Karel JMH, Wijnands E, et al. Integrating multiplex immunofluorescent and mass spectrometry imaging to map myeloid heterogeneity in its metabolic and cellular context. Cell Metab. 2022;34(8):1214-25.e6.

52. Boyle JJ, Johns M, Kampfer T, Nguyen AT, Game L, Schaer DJ, et al. Activating Transcription Factor 1 Directs Mhem Atheroprotective Macrophages Through Coordinated Iron Handling and Foam Cell Protection. Circulation Research. 2012;110(1):20–33.

53. Pang Z, Lu Y, Zhou G, Hui F, Xu L, Viau C, et al. MetaboAnalyst 6.0: towards a unified platform for metabolomics data processing, analysis and interpretation. Nucleic Acids Research. 2024;52(W1):W398–W406.

54. Xia W, Hilgenbrink AR, Matteson EL, Lockwood MB, Cheng J-X, Low PS. A functional folate receptor is induced during macrophage activation and can be used to target drugs to activated macrophages. Blood. 2009;113(2):438–46.

55. Ang O, Lim HY, Lim SY, Lau J, Azhar SHM, Thiam CH, et al. Loss of arterial resident LYVE-1 expressing macrophages exacerbates atherosclerosis. Atherosclerosis. 2023;379:S9.

56. Lim HY, Lim SY, Tan CK, Thiam CH, Goh CC, Carbajo D, et al. Hyaluronan Receptor LYVE-1-Expressing Macrophages Maintain Arterial Tone through Hyaluronan-Mediated Regulation of Smooth Muscle Cell Collagen. Immunity. 2018;49(2):326-41.e7.

57. Stöger JL, Gijbels MJ, van der Velden S, Manca M, van der Loos CM, Biessen EA, et al. Distribution of macrophage polarization markers in human atherosclerosis. Atherosclerosis. 2012;225(2):461–8.

58. Wu H, Gower RM, Wang H, Perrard X-YD, Ma R, Bullard DC, et al. Functional Role of CD11c+ Monocytes in Atherogenesis Associated With Hypercholesterolemia. Circulation. 2009;119(20):2708–17.

59. Hu X, Cai X, Ma R, Fu W, Zhang C, Du X. Iron-load exacerbates the severity of atherosclerosis via inducing inflammation and enhancing the glycolysis in macrophages. Journal of Cellular Physiology. 2019;234(10):18792–800.

60. Zheng K, Yang W, Wang S, Sun M, Jin Z, Zhang W, et al. Identification of immune infiltration-related biomarkers in carotid atherosclerotic plaques. Scientific Reports. 2023;13(1):14153.

61. Watanabe R, Sato Y, Ozawa N, Takahashi Y, Koba S, Watanabe T. Emerging Roles of Tumor Necrosis Factor-Stimulated Gene-6 in the Pathophysiology and Treatment of Atherosclerosis. International Journal of Molecular Sciences [Internet]. 2018; 19(2).

62. Zamani M, Cheng YH, Charbonier F, Gupta VK, Mayer AT, Trevino AE, et al. Single-Cell Transcriptomic Census of Endothelial Changes Induced by Matrix Stiffness and the Association with Atherosclerosis. Adv Funct Mater. 2022;32(47).

63. Slenders L, Wesseling M, Wei S, Boltjes A, Kapteijn DMC, van de Kraak P, et al. Endothelial-to-mesenchymal transition gene signature derived from single-cell transcriptomics of human atherosclerotic tissue associates with stable plaque histological characteristics. Vascular Pharmacology. 2025;159:107498.

64. Samaniego R, Palacios BS, Domiguez-Soto Á, Vidal C, Salas A, Matsuyama T, et al. Macrophage uptake and accumulation of folates are polarization-dependent in vitro and in vivo and are regulated by activin A. Journal of Leukocyte Biology. 2014;95(5):797–808.

65. Antohe F, Radulescu L, Puchianu E, Kennedy MD, Low PS, Simionescu M. Increased uptake of folate conjugates by activated macrophages in experimental hyperlipemia. Cell Tissue Res. 2005;320(2):277–85.

66. Jager NA, Teteloshvili N, Zeebregts CJ, Westra J, Bijl M. Macrophage folate receptor-β (FR-β) expression in auto-immune inflammatory rheumatic diseases: A forthcoming marker for cardiovascular risk? Autoimmunity Reviews. 2012;11(9):621–6.

67. Winkel LC, Groen HC, van Thiel BS, Müller C, van der Steen AF, Wentzel JJ, et al. Folate receptor–targeted single-photon emission computed tomography/computed tomography to detect activated macrophages in atherosclerosis: can it distinguish vulnerable from stable atherosclerotic plaques? Mol Imaging. 2014;13.

68. Gianfrancesco MA, Dehairs J, L’Homme L, Herinckx G, Esser N, Jansen O, et al. Saturated fatty acids induce NLRP3 activation in human macrophages through K+ efflux resulting from phospholipid saturation and Na, K-ATPase disruption. Biochimica et Biophysica Acta (BBA) - Molecular and Cell Biology of Lipids. 2019;1864(7):1017–30.

69. Soflaee MH, Kesavan R, Sahu U, Tasdogan A, Villa E, Djabari Z, et al. Purine nucleotide depletion prompts cell migration by stimulating the serine synthesis pathway. Nature Communications. 2022;13(1):2698.

70. Du X, Yang Y, Zhan X, Huang Y, Fu Y, Zhang Z, et al. Vitamin B6 prevents excessive inflammation by reducing accumulation of sphingosine-1-phosphate in a sphingosine-1-phosphate lyase–dependent manner. Journal of Cellular and Molecular Medicine. 2020;24(22):13129–38.

71. Endo N, Nishiyama K, Otsuka A, Kanouchi H, Taga M, Oka T. Antioxidant activity of vitamin B6 delays homocysteine-induced atherosclerosis in rats. Br J Nutr. 2006;95(6):1088–93.

72. Zhang P, Tsuchiya K, Kinoshita T, Kushiyama H, Suidasari S, Hatakeyama M, et al. Vitamin B6 Prevents IL-1β Protein Production by Inhibiting NLRP3 Inflammasome Activation. J Biol Chem. 2016;291(47):24517–27.

73. Zhao N, Huang Y, Wang YH, Muir RK, Chen YC, Wei J, et al. Ferronostics: Measuring Tumoral Ferrous Iron with PET to Predict Sensitivity to Iron-Targeted Cancer Therapies. J Nucl Med. 2021;62(7):949–55.

74. Raman SV, Winner MW, 3rd, Tran T, Velayutham M, Simonetti OP, Baker PB, et al. In vivo atherosclerotic plaque characterization using magnetic susceptibility distinguishes symptom-producing plaques. JACC Cardiovasc Imaging. 2008;1(1):49–57.

75. Vinchi F, Muckenthaler MU, Da Silva MC, Balla G, Balla J, Jeney V. Atherogenesis and iron: from epidemiology to cellular level. Front Pharmacol. 2014;5:94.

76. Sullivan JL. Iron and the sex difference in heart disease risk. Lancet. 1981;1(8233):1293–4.

77. Marques VB, Leal MAS, Mageski JGA, Fidelis HG, Nogueira BV, Vasquez EC, et al. Chronic iron overload intensifies atherosclerosis in apolipoprotein E deficient mice: Role of oxidative stress and endothelial dysfunction. Life Sci. 2019;233:116702.

78. Ma J, Ma HM, Shen MQ, Wang YY, Bao YX, Liu Y, et al. The Role of Iron in Atherosclerosis in Apolipoprotein E Deficient Mice. Front Cardiovasc Med. 2022;9:857933.

79. Kautz L, Gabayan V, Wang X, Wu J, Onwuzurike J, Jung G, et al. Testing the iron hypothesis in a mouse model of atherosclerosis. Cell Rep. 2013;5(5):1436–42.

